# Biased synaptic activation of dentate granule cells by exercise reflects inputs from the lateral entorhinal cortex

**DOI:** 10.64898/2026.02.22.707119

**Authors:** C Chatzi, AN Simmonds, A Veshagh, A Ellingson, M Krush, T McLean, E Schnell, GL Westbrook

**Author notes:** Neuroscience graduate program, Baylor College of Medicine (AN Simmonds). OHSU School of Medicine.

## Abstract

Hippocampal dentate granule cells receive multisensory information from the entorhinal cortex in a laminated and functionally segregated manner. We previously reported that brief periods of voluntary exercise in mice increased EPSCs and dendritic spines for inputs from the lateral, but not the medial, entorhinal cortex. Here we asked whether laminar specificity was due to molecular changes specific to distal granule cell dendrites or rather was dependent on upstream drive from the entorhinal cortex. Selective chemogenetic stimulation of either lateral entorhinal cortex (LEC) or the medial entorhinal cortex (MEC) increased granule cell dendritic spine density in the selected pathway. However, the preponderance of exercise-activated cells originated from LEC based on expression of an activity-dependent retrograde virus in Fos-TRAP mice. Our results indicate that the preferential activation by exercise reflects the drive of locomotor-related inputs from the lateral entorhinal cortex rather than selective molecular mechanisms in distal dendrites of dentate granule cells. How this activation pattern affects other salient stimuli involving contextual or spatial cues may underlie the benefits of exercise on learning and memory.

## INTRODUCTION

Neurons have the remarkable ability to process and respond to complex stimuli such as physical exercise and changes in an organism’s external environment. In rodents and humans, exercise/locomotion can improve learning and memory, likely due to an increase in hippocampal synaptic plasticity (van Praag et al., 1999; Dao et al., 2015; O’Callaghan et al., 2009; Vaynman et al., 2004; Brann et al., 2021). The dentate gyrus is particularly important in this regard as it receives diverse informational input from other brain regions and is involved in episodic memory, spatial navigation and the detection of novel stimuli (Squire, et al., 2004; Witter and Amaral, 2004; Amaral et al., 2007; Hainmueller and Bartos, 2020; Knierim et al., 2014). The medial entorhinal cortex is considered to convey spatial information via the medial perforant path (MPP), and the lateral entorhinal cortex codes for contextual and time cues via the lateral perforant path (LPP) that provides for pattern separation of incoming information (Witter and Amaral, 2004; Knierim et al., 2014; Yassa and Stark, 2011; Reagh and Yassa, 2014).

However, understanding which cells in the brain are activated by exercise, and how the preexisting circuitry is refined by this stimulus, has been difficult to examine. Likewise, most studies have used only periods of sustained exercise (van Praag et al., 1999; Farmer et al., 2004; Vaynman et al., 2004), making it more difficult to directly link exercise to specific functions at the synaptic and circuit level. In prior work in *eLife*, we used conditional Fos-TRAP mice to track the effects of a single exposure to exercise by permanently marking exercise-activated dentate granule cells in the hippocampus (Chatzi et al., 2019). This approach allows examination of the cascade of changes at the single neuron and circuit level post-exercise. This work surprisingly found that exercise selectively increased EPSCs and dendritic spines in the outer molecular layer inputs to granule cells and that expression of the BAR-domain containing protein Mtss1L (Metastasis suppressor 1-like) was necessary for this structural plasticity (Chatzi et al., 2019; Bruel-Jungerman et al., 2009).

Here we examined whether laminar-specific activation of dentate granule cells, induced by brief periods of voluntary exercise, resulted from synapse-specific molecular plasticity in the outer molecular layer, or reflected the relative balance of neural activity originating from the entorhinal cortex. The MEC and LEC selectively project via the medial and lateral perforant paths to the middle and outer molecular layers of the dentate gyrus, respectively (Witter and Amaral, 2004; Knierim et al., 2014). Our results indicate that neural activity from the lateral entorhinal cortex underlies the increase in granule cell activity and selectively increased structural plasticity in the outer molecular layer. As these inputs have been implicated in novelty detection (Hainmueller and Bartos, 2020; Knierim et al., 2014; Witter and Amaral, 2004), our results provide further support for a role of exercise in learning and memory.

## RESULTS

### Laminar-specific activation of dentate granule cells by entorhinal DREADDs

Incoming stimuli preferentially activate discrete layers on the distal dendrites of granule cells (Chatzi et al., 2019). For example, single bouts of voluntary exercise (2 hrs) transiently increase EPSCs and dendritic spines in the outer molecular layer (OML) (Chatzi et al., 2019). To investigate the differential contribution of the entorhinal cortex subregions to the activation and plasticity of dentate granule cells, we took advantage of the spatial segregation of the lateral or medial entorhinal cortex (LEC or MEC) fiber pathways, which target the middle and outer molecular layers of the dentate gyrus respectively (Witter and Amaral, 2004; Brun et al., 2002). First, we used a chemogenetic approach to selectively activate inputs from LEC or MEC. Because dentate granule cells receive direct EC inputs from neurons in layer 2a of the EC, we injected wild-type mice unilaterally with adeno-associated virus (AAV) in either the superficial layer of LEC or MEC to express the excitatory DREADD, hM3D(Gq)-mCherry. (Alexander et al., 2009). Administration of CNO (i.p.) greatly increased c-Fos^+^ granule cells on the ipsilateral dentate gyrus, compared to the contralateral side (CNO-only vs. LEC DREADDs+CNO, n=4, p=0.003; CNO-only MEC vs. MEC DREADDs+CNO, n=4, p=0.003, Figure 1A).

**Figure 1:**
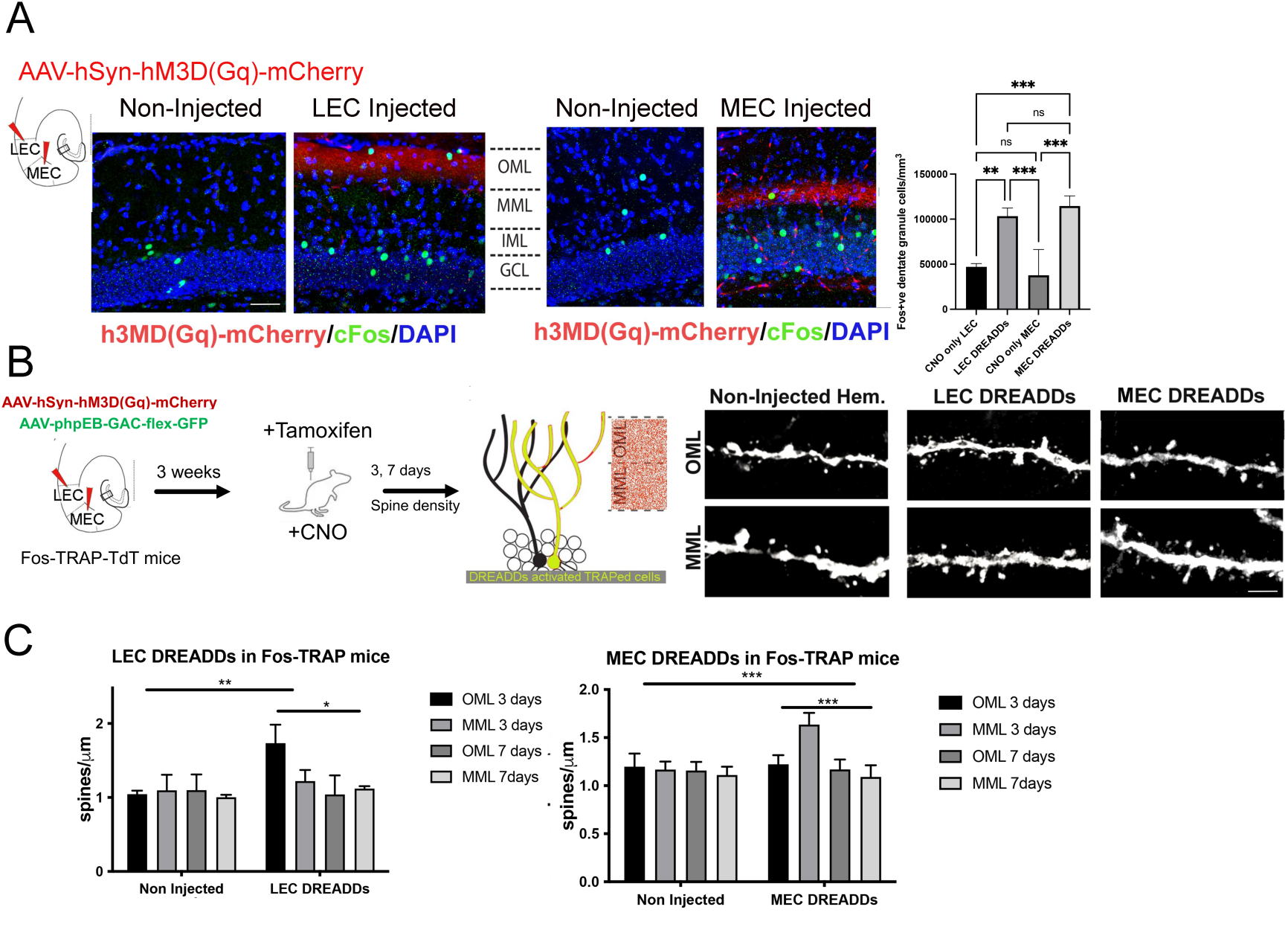
A pharmacogenetic approach effectively activates either the lateral and medial perforant path. A. An AAV was injected unilaterally to express the excitatory DREADD hM3D(Gq) in either the lateral (LEC) or medial entorhinal cortex (MEC). Three weeks later, mice were injected with CNO (1mg/kg, i.p.) and sacrificed 2hrs later to examine c-FOS expression in dentate granule cells. Dentate gyrus coronal sections show OML or MML fibers selectively expressing hM3D(Gq)-mCherry (red band) after LEC (left) or MEC (right) injections. CNO-mediated chemogenetic activation of either MEC or LEC increased c-FOS^+^ (green) granule cell density. c-FOS^+^ dentate granule cell counts were quantified in CNO only and CNO^+^ excitatory DREADD (LEC and MEC). c-Fos^+^ granule cells were significantly higher in the hemisphere injected with excitatory DREADD hM3Dq in either the LEC or MEC compared to CNO only treated contralateral hemisphere. (CNO Only LEC: 47106±3560, n=3, LEC DREADDs: 103363± 8990, n=4, CNO Only MEC: 37708± 28545, n=4, LEC DREADDs: 114536± 11237, n=4, ANOVA with Tukey’s multiple comparisons. Symbols represent: *** *p* < 0.001). Scale bar: 50 µm B. Experimental timeline for chemogenetic excitation of either the lateral or medial perforant path in Fos-TRAP mice for quantification of dendritic spines. Fos-TRAP:TdTomato mice received unilateral stereotaxic injections of AAV-hM3D-mCherry into either the MEC or LEC, and AAVphpEB-CAG-flex-GFP retroorbitally. Three weeks later, tamoxifen was injected (150mg/kg, i.p.) 24hrs before CNO administration (1mg/kg, i.p.) to label DREADD TRAPed granule cells with both tdTomato and GFP. Mice were sacrificed 3 or 7 days after CNO injection. Representative images are shown of LEC and MEC DREADD TRAPed granule cell dendrites (tdT signal) in the OML and MML from mice in the non-injected (CNO only) or injected (DREADDs+CNO) hemisphere. Scale bar = 2 µm. C. Spine densities were significantly increased in the OML of LEC DREADD TRAPed granule cells 3 days post-CNO (two-way ANOVA, Non-Injected OML 3 days: 1.04 ± 0.02, LEC DREADDs OML 1.73 ± 0.11, *p* < 0.01, n = 5), while spine density in the MML was unaffected (two-way ANOVA, no interaction between non-Injected and LEC DREADDs groups, Non-Injected MML 3 days: 1.10 ± 0.13, LEC DREADDs MML 1.22 ± 0.08, p < 0.05, n = 5). Similarly, spine densities were significantly increased in the MML but not OML of MEC DREADD TRAPed granule cells 3 days post CNO injection (two-way ANOVA, Non-Injected MML 3 days: 1.16 ± 0.08, MEC DREADDs MML 1.63 ± 0.12, *p* < 0.01, n = 5; Non-Injected OML 3 days: 1.15 ± 0.09, MEC DREADDs OML 1.17 ± 0.10, *p* < 0.05, n = 5). Spine densities of TRAPed dentate granule cells returned to baseline in both OML and MML 7 days after chemogenetic activation of either the LEC or the MEC (no interaction *p* < 0.05, n = 5 animals per experimental condition). Graphs display mean ± standard deviation.

To visualize individual dendritic segments, we injected AAVphpEB-CAG-flex-GFP retroorbitally to achieve sparse labeling of activated granule cells. DREADD activation in LEC caused a 50% increase in granule cell spine density at 3 days in the OML, but not the MML (Non-injected:OML 3 days vs. LEC DREADDs OML 3 days, n=5, p< 0.01; Non-injected:OML 3 days vs. LEC DREADDs: MML 3 days, n=5, p=0.148, Figure 1B, 1C, left). Similarly, DREADD activation of inputs from MEC increased spine density in the MML with no change in the OML (Non-injected:MML 3 days vs. MEC DREADDs MML 3 days, n=5, p< 0.05; Non-injected:MML 3 days vs. MEC DREADDs: OML 3 days, n=5, p=0.267, Figure 1B, 1C, right). Spine densities returned to baseline levels by 7 days post-chemogenetic activation in both layers (Figure 1C), further supporting a transient response to a single physiological stimulus, as previously reported (Chatzi et al., 2019). These results indicate that input from either the LEC or MEC is capable of activating granule cells and increasing dendritic spines. Thus, the preferential activation of distal dendrites in the OML by voluntary exercise must involve additional mechanisms.

### Laminar-specific activation by exercise does not involve newly-generated granule cells

We and others have shown that adult-born dentate granule cells at 3-4 weeks post-mitosis receive substantial functional input from the LEC, but only weak input from the MEC (Vivar et al., 2012; Woods et al., 2018; Gu et al., 2012). To investigate whether the OML-specific increase in spine density following exercise could be attributed to the preferential activation of adult-born granule cells (Schmidt-Hieber et al, 2004; Sahay et al., 2011; Marlatt et al, 2012; Vivar and van Praag, 2013; Vivar et al., 2016), adult Fos-TRAP mice underwent a single bout (2 hours) of exercise, which causes a 3-5 fold increase in activated granule cells (Chatzi et al., 2019). Three days later, we examined co-labeling of exercise-TRAPed cells with Doublecortin/DCX, which labels cells at approximately 7-21 days post-motosis), and with Calbindin to label mature granule cells (Zhao et al., 2006). Co-labeled exercise-TRAPed cells exhibited highly arborized dendritic processes extending fully across the molecular layer, consistent with mature granule cell morphologies (Figure S1 A). However, less than 2% of exercise-TRAPed granule cells were DCX^+^ (Calbindin⁺/Exercise-TRAPed⁺ overlap: 72.4 ± 5.2%, DCX⁺/ Exercise-TRAPed overlap: 1.2% ± 0.01%, n=5, p < 0.0001, Figure S1A, right), indicating that exercise-TRAPed cells are predominately mature granule cells.

Newborn neurons begin to receive functional excitatory inputs from the entorhinal cortex at 3 weeks post-mitosis (Ge et al., 2007), but reach stable dendritic spine densities and synaptic properties by 6 weeks post-mitosis, at which point they are generally regarded as mature granule cells (Ge et al., 2008; Woods et al., 2018). As an alternative method to determine whether exercise-TRAPed cells represent an immature cohort, we examined the birthdate of exercise-TRAPed cells using the mitotic marker 5-bromo-2’-deoxyuridine (BrdU) to label cells born either 3 or 4 weeks before exercise. There was negligible co-labeling of BrdU in exercise-TRAP^+^ cells, indicating that exercise-TRAPed cells represent mature granule cells (21 days post BrdU, % BrdU+exercise-TRAPed^+^ 0.02%± 0.09, n=3; 28 days post BrdU, % BrdU+exercise-TRAPed^+^ 0.04± 0.08, n=3; Figure S1B).

### Functional entorhinal input using an activity-dependent retrograde AAV

The OML and MML receive dense axonal inputs from LEC and MEC, respectively (Witter and Amaral, 2004). To obtain a direct measure of which entorhinal afferents drive activity onto exercise-TRAPed cells, we used an activity-dependent retrograde tracing approach in which viral particles are incorporated into activated presynaptic terminals (Guenthner et al., 2013; Cai et al., 2016). We injected Cre-dependent retrograde AAV (AAVrg-flex-GFP) into the terminal fields of the perforant pathway in both OML and MML of Fos-TRAP mice. Exercise labeled AAVrg-flex-GFP+ axons in the OML and MML upstream of exercise-TRAPed dentate granule cells, indicating that exercise induced axonal labeling in inputs from both LEC and MEC (Figure 2A, bottom left panel). There was also some AAVrg-flex-GFP+ axons in the molecular layer in the home cage condition (Figure 2A, bottom right panel), reflecting the ongoing baseline activity of granule cells. Interestingly there was no labeling of axons in the inner molecular layer (IML), which arise from hilar mossy cells, but robust labeling in the supragranular layer, possibly representing inputs from the supramammillary nucleus (Pan and McNaughton, 2004; Hashimotodani et al., 2018).

**Figure 2.**
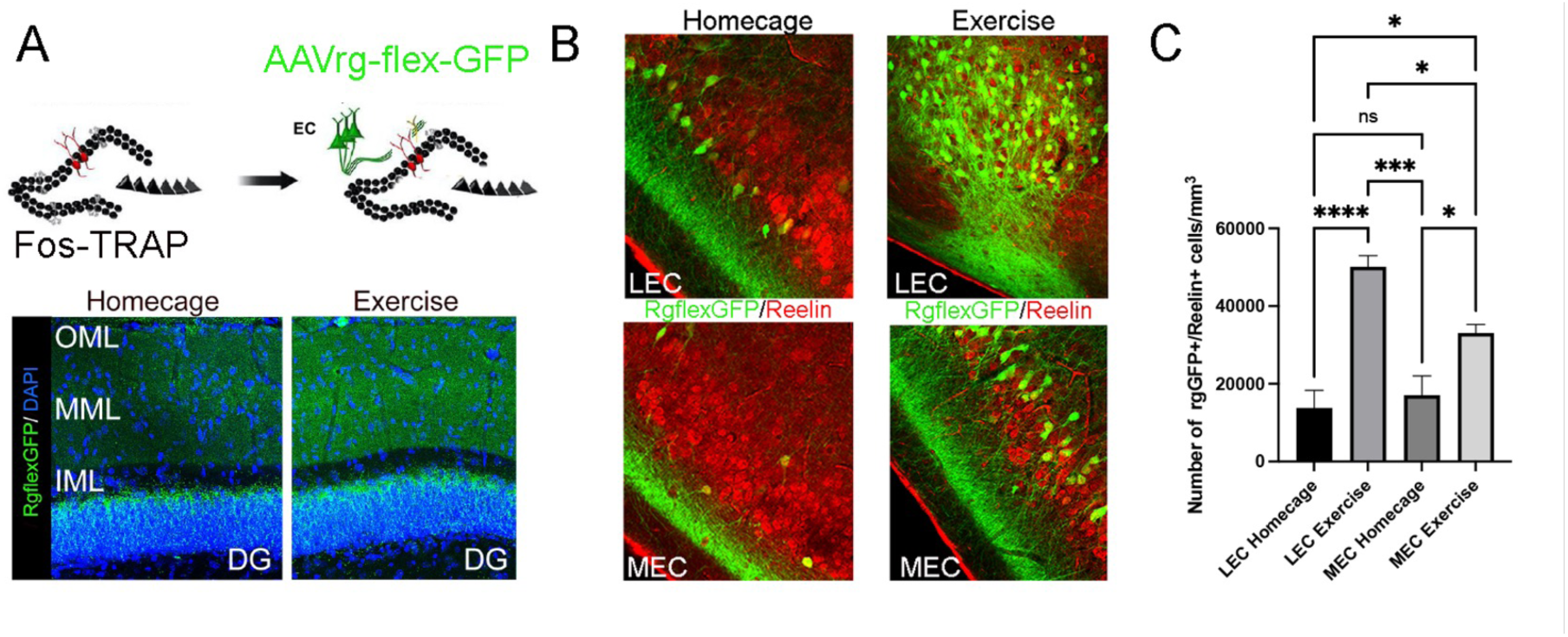
Active perforant path inputs to dorsal exercise-TRAPed dentate granule cells from the entorhinal area originate largely from the LEC. A. (Top) Schematic of the experimental paradigm examining the perforant path connectome upstream of exercise-activated neurons. A Cre-dependent retrograde AAV was stereotactically injected into perforant pathway terminal fields (OML and MML) in the dentate gyrus of Fos-TRAP mice. Three weeks later, tamoxifen administration labelled exercise-dependent presynaptic neurons in the EC projecting to the dentate gyrus. (Bottom) Representative images of lateral and medial entorhinal cortex axons labelled in an activity-dependent manner by Cre-dependent retrograde AAV expressing GFP, injected into the OML and MML of the dentate gyrus of Fos-TRAP mice. Exercise increased the fluorescence intensity of AAVrg-flex-GFP along the middle and outer molecular layer compared to home cage controls. B. Immunolabelling of layer IIa Fan cells with Reelin (red) and Fos-TRAP: retrograde-GFP^+^ in LEC and MEC (left) revealed double labelling of exercise-activated cells that project to the dentate gyrus C. The density of double labelled Fos-TRAP: retrograde-GFP^+^ and Reelin^+^ neurons was compared between LEC and MEC in home cage and exercised Fos-TRAP mice (LEC Home cage:13557±7893, n=3, LEC Exercise 50100±6409, n=5, MEC Home cage, 17088±9830, n=4, MEC Exercise: 33093±4390, n=4). Exercised mice showed a significantly larger density of Reelin^+^ Fos-TRAP: retrograde-GFP^+^ cells than mice in home cage conditions in both the LEC and the MEC. However, the number of rgGFP^+^/Reelin^+^ cell density in LEC was significantly greater than the MEC. LEC Exercise n=5, MEC Exercise n=4, p=0.02 Two-way ANOVA with multiple comparisons. Scale bars: 100 µm (DG overview), 50 µm (LEC/MEC)

To examine the source of exercise-activated entorhinal inputs, we assessed entorhinal cells that were retrograde-GFP^+^ labeled. Exercise increased the number of labeled cells in both MEC and LEC compared to home cage. However, there were more double-labeled cells in LEC compared to MEC (Figure 2B, C). Double staining with Reelin, a marker of LEC layer 2, DG-projecting entorhinal cells, confirmed that the number of Fos-TRAP: retrograde-GFP^+^ cells was higher in LEC (LEC Exercise n=5, MEC Exercise n=4, p=0.02, Figure 2C). Thus, both LEC and MEC excite exercise-TRAPed cells through direct projection to the dentate gyrus, but there was a 2-fold greater number of activated LEC neurons that project to exercise-activated granule cells compared with MEC.

### Activation of cells in LEC by exercise is stochastic

Our prior results indicated that exercise activation of dentate granule cells was stochastic because distinct populations of granule cells were activated by two identical bouts of exercise separated by 3 days (Chatzi et al., 2019). We thus asked whether exercise-activated cells in the lateral entorhinal cortex (Fos-TRAP^+^) specifically respond to exercise or rather reflect stochastic activation of cells in layer II, the main source of inputs to the dentate gyrus. We compared exercise-TRAPed (Fos-TRAP^+^) cells in the LEC following an initial bout of exercise with Fos^+^ <u>immunolabeled</u> cells activated by a second bout of exercise 3 days later (Figure 3A, B). To assess the number of activated cells, we calculated the fraction of labeled cells compared to the total number of DAPI-labeled cells counted (not shown) in LEC layer 2. Strikingly, the merge panel In Figure 3B showed an extremely low level of overlap (<0.01% that did not differ from the overlap expected by multiplying the percentage labeling from the two bouts of exercise, i.e. random chance; n=4, p=0.91). Thus, the strong exercise activation of cells in LEC does not reflect stimulus (exercise)-specific cells in the entorhinal cortex.

**Figure 3:**
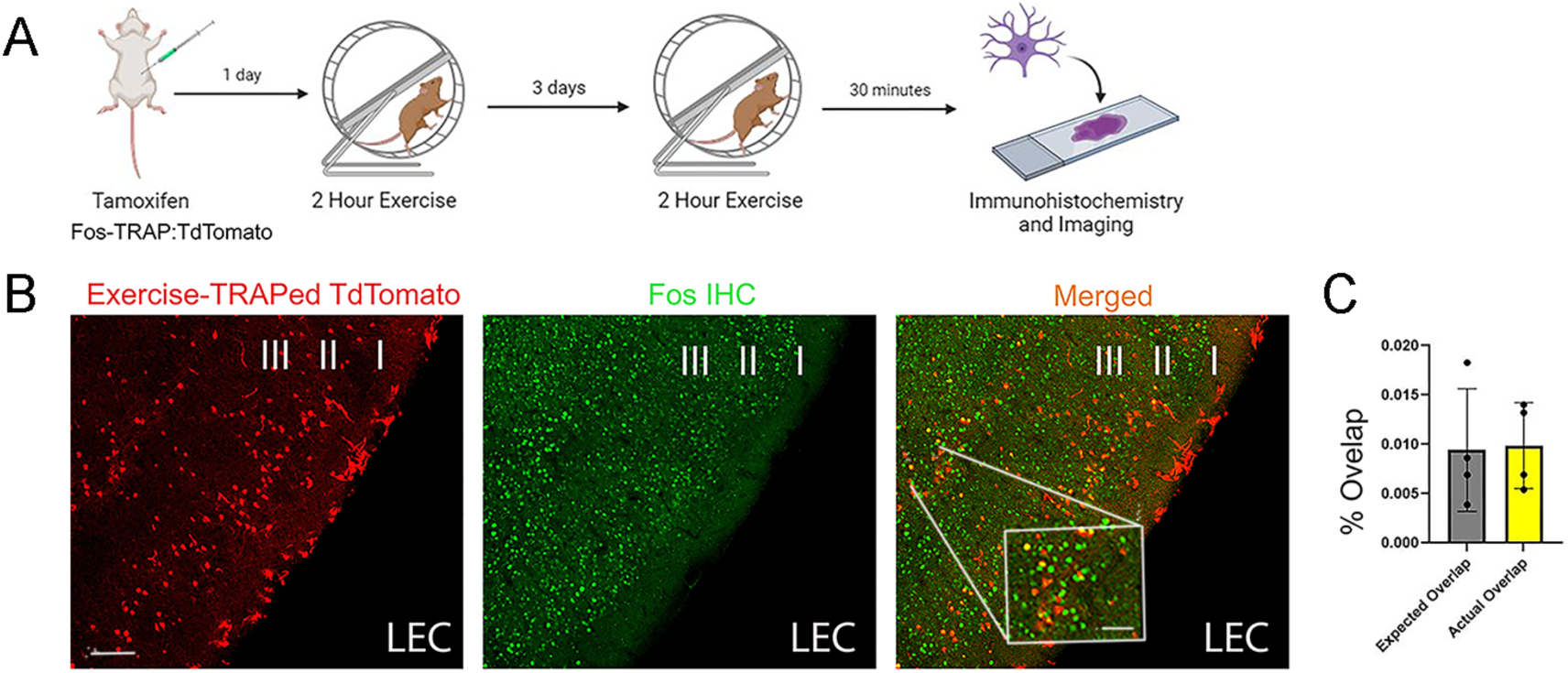
Stochastic Activation of Cells in the LEC. A. Experimental timeline for labeling of activated granule cells in mice during two discrete rounds of exercise separated by 3 days. Fos-TRAP mice were given intraperitoneal tamoxifen (150 mg/kg) 24 hours before the first two-hour exercise session to label activated granule cells. Following this session, mice were returned to their home cage. Three days later, mice were re-exposed to the running wheel for an additional two hours and perfused 30 minutes later. Lateral entorhinal cortex (LEC) sections were stained for tdTomato and c-Fos, respectively to label cells activated during the two sessions. B. Representative LEC images from double-exercised Fos-TRAP mice, showing fos-TRAP^+^ cells in red, c-Fos+ cells in green. The merged image highlights overlapping cells in yellow in the LEC. Scale bars: 100 µm.Inset, 20 µm. C. Histogram showing the percentage of cells labeled with fos-TRAP and c-Fos-labeled cells in LEC layer II. The observed overlap (yellow: 0.01±0.005, n=4) did not differ from the expected overlap due to chance based on the fraction of labelled cells (gray: 0.009±0.007, n=4), p= 0.9080). See text for how percentage was calculated.

### Silencing of perforant path inputs with tetanus toxin light chain

Despite the preferential activation of granule cells by inputs from the LEC in exercise, our results indicate that activation of MEC afferents via DREADDs can also drive structural plasticity of Fos-TRAPed dentate granule cells, and that retrogradely labeled MEC neurons in layer 2 can be exercise-activated. To examine the extent of the biased activation, we used viral expression of GFP-tagged tetanus toxin light chain (TeLC-GFP; Murray et al., 2011). To first validate the efficacy of this tetanus light chain vector, we injected one LEC hemisphere with a viral control GFP and the contralateral LEC hemisphere with TeLC-GFP virus. After 3 weeks, we recorded extracellular field potentials in an acute brain slice evoked by paired electrical stimulation of either lateral perforant path (LPP) or medial perforant path (MPP). Field potentials evoked by paired LPP stimulation were robust in the control hemisphere (CtrlGFP-LEC, Figure 4A, left, upper row), but were completely eliminated in the LPP of the TeLC-expressing hemisphere (TeLC-LEC) (Figure 4A, right, upper row). In contrast, stimulation of the MPP was robust in both hemispheres (Figure 4A, lower row). Paired pulse depression was greater for the MPP than LPP stimulation (P_2_/P_1_: LPP =0.88, MPP=0.75 in the example shown, Figure 4A), consistent with prior reports in rodent dentate gyrus (Colino and Malenka, 1993; Asztely et al., 2000), and confirming the accuracy of the placement of stimulating electrodes.

**Figure 4:**
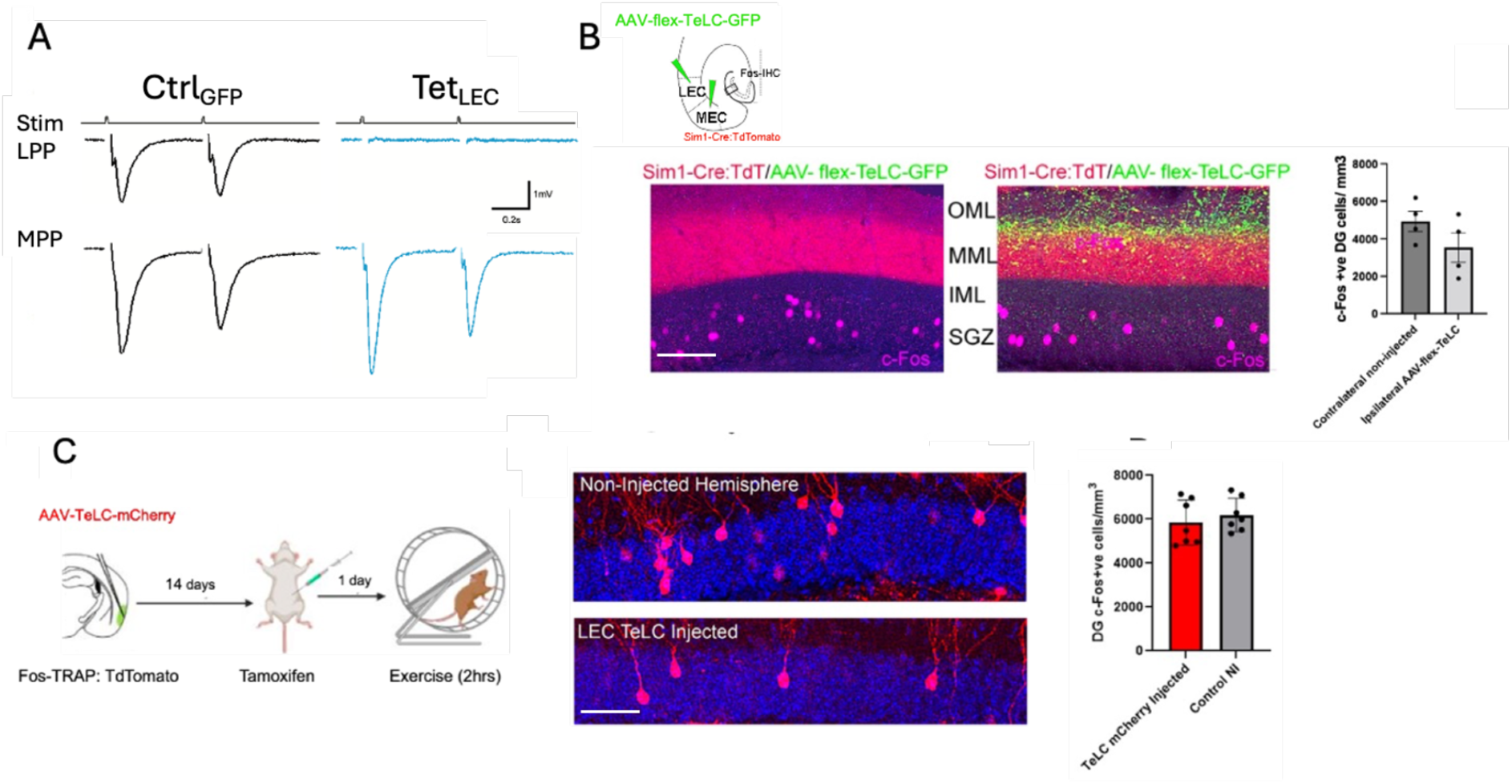
Functional silencing of LEC inputs is not sufficient to block exercise-induced granule cell activation. A. To validate the TeLC-GFP viral construct, the LEC in one hemisphere was injected with AAV- flex-TeLC-GFP and the contralateral LEC was injected with a control AAV- flex -GFP virus. Extracellular field recordings in acute brain slices were made in the molecular layer of the dentate gyrus following selective stimulation of either the lateral perforant path (LPP, top row) or medial perforant path (MPP, bottom row). Paired stimuli with a bipolar electrode revealed field EPSPs following LPP or MPP stimulation with the control GFP virus (left column, CtrlGFP); however, the field EPSPs following injection with TeLC-GFP showed complete block following LPP stimulation (right column, TeLC, consistent with silencing of the LPP. B. Schematic representation of AAV-flex-TeLC-GFP injection in Sim1-Cre-:TdT mice, targeting the right hemisphere LEC <u>and</u> MEC layer IIa (upper). Representative dentate gyrus images show c-Fos^+^ labeling in magenta and axons expressing AAV- flex -TeLC-GFP^+^ in green (left). There was not a significant difference in c-Fos^+^ (activated) dentate granule cells between the injected (right image) and non-injected hemispheres (left image). Scale bar = 100 µm. C. Timeline for Fos-TRAP activation following a unilateral AAV5-TeLC-mCherry injection in LEC layer IIa of Fos-TRAP mice. Mice received tamoxifen 24 hours before the experiment, followed by two hours of voluntary wheel running and perfusion three days later. Representative images of c-Fos-activated cells in the dentate gyrus of TeLC-injected (lower) and non-injected (upper) hemispheres. c-Fos^+^ cells (red); DAPI (blue). There was no significant difference in the number of c-Fos^+^ DGCs between the AAV5-TeLC-mCherry-injected and non-injected hemispheres (AAV5-TeLC mCherry-injected: 5858 ± 1026 /mm^3^, Non-injected: 6198 ± 761 /mm^3^, p=0.5, n=7). Scale bar = 20 µm.

We then targeted layer 2 of <u>both</u> LEC and MEC with AAV flex TeLC-GFP using Sim1-Cre mice, which selectively labels Fan cells in layer 2 of the entorhinal cortex (Sürmeli et al., 2015; Vandrey et al., 2020). After 3 weeks for expression of virus, GFP^+^ perforant path axons were present throughout the MML and OML, consistent with labeling of perforant path axons from Sim1-expressing cells in the entorhinal cortex (Figure 4B). We compared total neuronal activity (by Fos immunohistochemistry) in dentate granule cells following unilateral TeLC-GFP injection of <u>both</u> LEC *and* MEC compared to the non-silenced contralateral hemisphere (Figure 4B,) Perhaps surprisingly, the number of c-Fos^+^ dentate granule cells in the TeLC-GFP-injected hemisphere was not significantly lower than the non-silenced hemisphere (Figure 4B, right), indicating that our silencing of EC afferents was not sufficient to alter the baseline level of activity of granule cells (Figure 4B, n=4, p=0.08). We also injected AAV-flex-TeLC-GFP in the superficial layers of the LEC and assessed the number of exercise-TRAPed granule cells in the dentate gyrus. There was no reduction in exercise-activated granule cells, presumably a result of continuing activity of inputs from the MEC (p=0.4974, n=7, Figure 4C).

### Pattern of granule cell activation following short-term environmental enrichment

To examine whether biased synaptic activation of dentate granule cells as determined by exercise-induced activity is specific to locomotion-induced activity, we examined the effects of short-term environmental enrichment (EE). Prior studies of EE in mice have used sustained stimuli including objects, running wheels and other mice over periods of several weeks (Kempermann et al., 1997; Eadie et al., 2005; Stranahan et al., 2007). In order to compare EE to our experiments with brief exercise, we used a modified EE environmental arena that included objects and tunnels but no running wheels to avoid confounding effects of locomotion (Figure 5A). Neural activity in the dentate gyrus of adult mice was assessed in the home cage after 6 or 24 hours of EE exposure using immunolabeling for the immediate early genes, Fos and Arc (Ramirez-Amaya et al., 2005; Guzowski et al., 2001). EE resulted in a nearly 2-fold increase in Fos^+^ cells (red) at 24hrs post EE (p=0.0009, n=4, Figure 5B). Arc (green) showed a similar increase but was not quantified (Figure 5B). Assessment of dendritic spines in the mice following 24 hours of EE exposure in the OML and MML were greater than previously observed in home cage mice (Chatzi et al., 2019), but did not reveal a significant difference in spine density between the layers (p= 0.144, n=4, Figure 5C).

**Figure 5.**
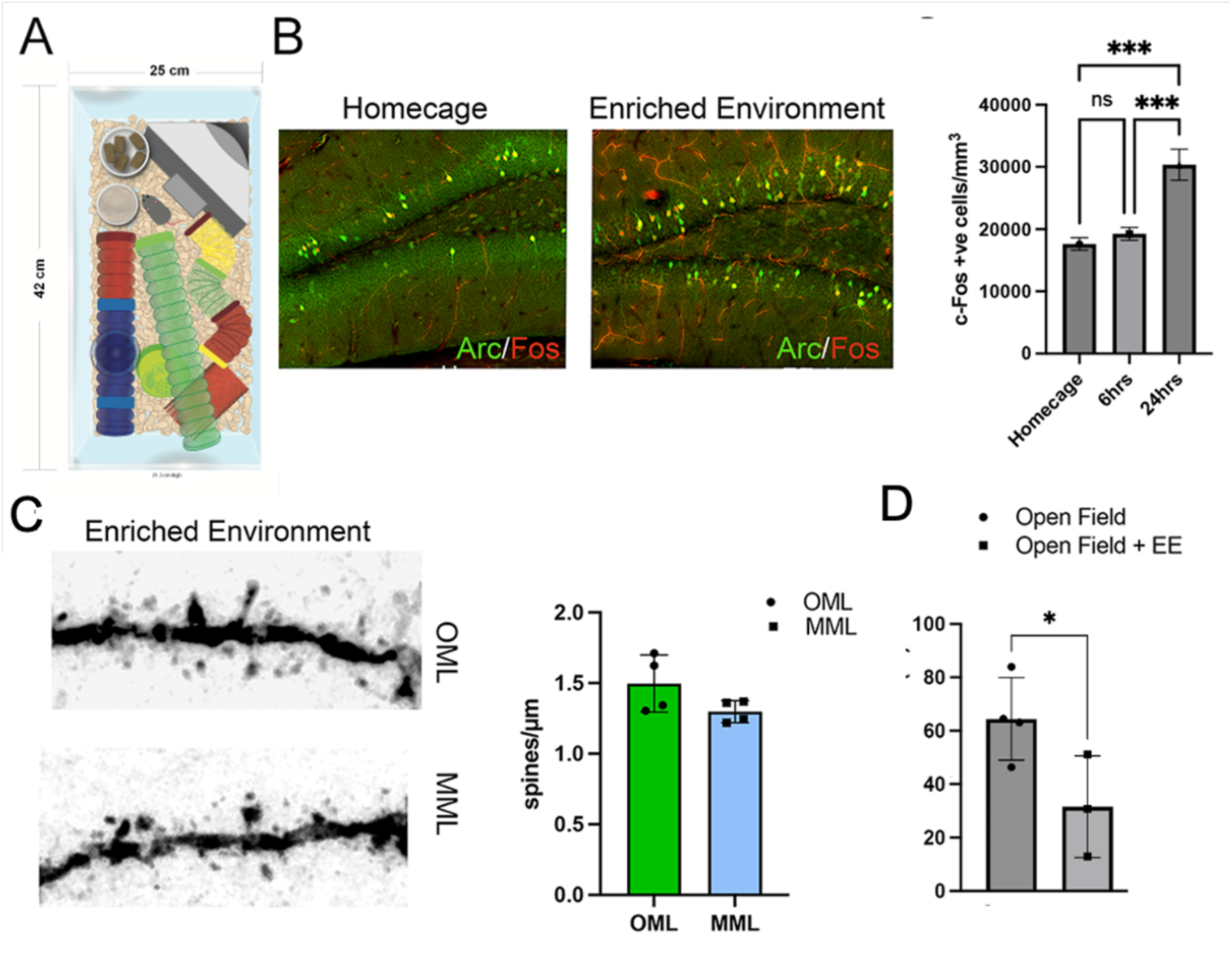
Short-term enriched environment (EE) increased neuronal activation, but did not alter the distribution of dendritic spines between MML and OML. A. A schematic of the objects used for EE including running wheels, tubes and various objects. B. Mice remained in home cage or were exposed to EE for 6 or 24 hours, then were assessed by immunohistochemistry by Fos (red) or Arc (green). Activated granule cells (Fos^+^) were increased after 24 hours of EE but not after 6 hours (home cage: 16878±2098, 6 hrs EE: 21867±5813, 24hrs EE: 35133±5647, cells/mm3, n=4). Scale bar = 100 µm. C. Dendritic spines in the middle and outer molecular layers of the dentate gyrus showed similar densities in mice exposed to EE for 24 hours (OML: 1.50 ± 0.20 spines/μm; MML: 1.30 ± 0.08, spines/μm, n=4). Scale bar = 2 µm. D. In a separate cohort of mice, which were acclimated to an open field arena, total locomotion in the arena was reduced by inclusion of the EE objects compared to empty open field (Open field: 64.5±8.8 meters, n=4, EE: 31.6± 9.1 meters, n=3).

To test whether the increased activation was simply a result of increased locomotion, in a separate experiment, we used open field arena to quantify distance traveled. Mice were brought to the arena on day 1 for acclimation, and then returned on day 2 for testing with or without EE objects. For the 20 minute test period, there was a decrease in the distance traveled in the enriched environment, which is likely explained by increased exploration of the novel objects (Open Field, n=4, Open Field+EE: n=3, p=0.008, Figure 5D).

## DISCUSSION

Our results provide further understanding of the circuit alterations that occur in response to voluntary exercise in the mouse dentate gyrus. The “locomotor drive” which activates granule cells and increases dendritic spines in a pathway-specific manner is dependent on the pattern of activation of entorhinal inputs upstream of the dentate gyrus, and thus could not be attributable to laminar-specific localization of Mtss1L (Chatzi et al, 2019), synapses in the outer molecular layer that receive inputs from that lateral entorhinal cortex (Witter and Amaral, 2004; Knierim et al., 2014). Activation from LEC relative to MEC was a bias rather than a hard distinction between contextual and spatiotemporal inputs attributable to those perforant path lamina. Thus, brief voluntary running in our experiments served as a relatively selective, but not absolute, drive from the lateral entorhinal cortex, and presumably downstream from exercise-triggered multimodal cortical inputs to LEC. However, activated cells in LEC and in the dentate gyrus were not “exercise-specific” as repeated inputs generated distinct cohorts of activated granule cells and EC projection cells, consistent with a relative stochastic cellular activation. However, changes in dendritic spines in exercise-activated cells reflect local neural activity, as exhibited by expresssion of Fos in exercise-activated cells (Chatzi et al., 2019).

### Comparison to prior results

Assessing neural circuit activity using c-FOS immunohistochemistry as well as the use of Fos-TRAP mice as a proxy for neural activity has been well-validated (West and Greenberg, 2011; Guenthner et al., 2013; Chatzi et al., 2019; Cappaert et al., 2015), but almost certainly requires action potential generation in individual neurons to provide a sufficient stimulus for fos activation in the nucleus (Bruel-Jungerman et al., 2009; Guzowski et al., 2001). This approach is well suited to the dentate gyrus in which neurons are relatively silent (Jung and McNaughton, 1993; Hainmueller and Bartos, 2020), such that the network shows only a few granule cells active at baseline, characteristic of a sparse network (Leutgeb et al., 2007; Senzai and Buzsáki, 2017), and consistent with a role of the dentate gyrus in pattern separation (Yassa and Stark, 2011; Sahay et al., 2011; Berron et al., 2016; Clelland et al., 2009; van Dijk and Fenton, 2018). This low basal activity is also consistent with the rare spontaneous calcium transients in granule cells at baseline (Danielson et al., 2016; Senzai and Buzsáki, 2017).

Several caveats do affect the interpretation of Fos^+^ cells in our experiments. The first is that each bout of exercise activated distinct populations of granule cells, suggesting that the stimulus was stochastic at a cellular level (Chatzi et al., 2019). This pattern also appears to apply to exercise-associated activation of layer II entorhinal cortex, which provides the primary perforant path input to the dentate gyrus. This result undermines the superficially attractive idea that neurons Fos-tagged by recent exercise activation are primed for subsequent salient inputs. In fact, there is evidence that another immediate early gene, deltaFosB, may suppress subsequent activation of Fos (McClung et al., 2004; Renthal et al., 2008; Nestler, 2015; Robison and Nestler, 2011). Fos^+^ granule cells in the dentate gyrus are also limited to mature cells, because newborn granule cells in the subgranular zone are not labeled with Fos^+^ and thus were not represented in our experiments (Gu et al., 2012; Woods et al., 2018; Allen Institute for Brain Science, 2008, https://mouse.brain-map.org/experiment/show/79912554). Although we previously observed that newborn granule cells, labeled with retrovirus and recorded at 3 weeks later, showed preferential synaptic inputs in the outer molecular layer (Woods et al., 2018), the biased exercise activation from LEC could not be attributed to newborn granule cells.

### What is special about running as a stimulus to the dentate gyrus?

We used brief periods of voluntary running as a “natural” physiological stimulus that would primarily engage CNS mechanisms, rather than systemic changes in other organ systems such as cardiovascular, muscle and liver which are associated with sustained periods of physical exercise (van Praag et al., 1999; Dao et al., 2015; Mandolesi et al., 2018; Lourenco et al., 2019; Mu et al., 2022). Our approach increased the density of Fos^+^ (exercise-activated) granule cells and specific changes in neuronal gene expression in activated cells within the dentate gyrus (Chatzi et al., 2019). The inverse BAR protein Mtss1L (Metastasis suppressor 1-like) was upregulated by exercise-induced neural activity, and necessary for the functional increase in EPSPs and dendritic spines in the OML, as these changes were blocked by shRNA-mediated knockdown of Mtss1L (Chatzi et al., 2019). However, our results here indicate that selective localization of Mtss1L at OML synapses would be insufficient to explain the effects of voluntary exercise. Likewise, we were unable to demonstrate OML synapse-specific expression of Mtss1L using either available antibodies or with a His-tagged KI mouse (data not shown). However, human mutations in Mtss1L (Mtss2) have also recently been reported as a cause of intellectual disability (Huang et al, 2022) indicating a need for further studies of the inverse BAR domain proteins in brain function (Chatzi et al., 2021).

### Why the LEC “bias”?

Generally, LEC is considered to process input from multiple cortical areas (Eichenbaum et al., 2007; Ranganath and Ritchey, 2012; Schultz et al., 2012; Knierim et al., 2014; Witter and Amaral, 2004; Gonzalez-Parra et al., 2025; Ritchey et al., 2015) with contextually relevant sensory information (Eichenbaum et al., 2007; Schultz et al., 2012). This information then converges on the dentate gyrus to shape the what/when/where aspects of an experience. The context of an experience must depend on multimodal sensory and motor inputs (Knierim et al., 2014; Schultz et al., 2012). However, we initially were surprised that brief periods of voluntary exercise would preferentially activate LEC inputs to the dentate gyrus. Because the amplitudes of evoked perforant path excitatory inputs to OML and MML are similar (Brun et al., 2002; Witter and Amaral, 2004; Leutgeb et al., 2007) as is the axonal density, it is the activity of these inputs that determines the bias, a result that was supported by our experiments with activity-dependent retrograde labeling of LEC and MEC neurons that target the dentate gyrus (Guenthner et al., 2013; Cai et al., 2016). Although the anatomical inputs to LEC and MEC are well described (Witter and Amaral, 2004; Cappaert et al., 2015), the specific pathways that functionally contribute under conditions of brief exercise will require further study.

### Locomotion and brain function

The impact of locomotion on cognition is epitomized by the well-acknowledged value of short-term exercise on learning and memory (van Praag et al., 1999; Mandolesi et al., 2018; Dao et al., 2015; Lourenco et al., 2019; Mu et al., 2022). However, the effects of exercise and locomotion are pleiotropic, involving multiple organ systems as well as the brain (Mandolesi et al., 2018; Voss et al., 2013). The beneficial effects of exercise on other organ systems are usually evaluated after continuous exposure over weeks to months, but the effects on learning can be seen in humans in as little as 24 hours (Mandolesi et al., 2018), suggesting that there are short term changes in brain function. Such short-term changes on the order of seconds to minutes to days are expected to involve changes at specific synapses as well as alterations in signaling that may involve cells and circuits not directly affected by exercise-induced changes in neural activity (Dao et al., 2015; Mu et al., 2022). Our results indicate that even brief periods of exercise drive neural activity and subsequent structural plasticity in the dentate gyrus (Chatzi et al., 2019; Bruel-Jungerman et al., 2009; Lourenco et al., 2019). Although these exercise-induced changes were confined to specific laminar inputs from the LEC, further work will be needed to explore whether these changes are exercise-specific or reflect the patterns of neural activity generated by locomotion compared to other stimuli. If one accepts that LEC inputs carry information relative to context, then locomotion is more than a simple motor stimulus, but rather an integrated sensorimotor input (Schultz et al., 2012; Knierim et al., 2014). Studies in the visual system have shown that locomotion alters the processing in the visual cortex (Niell and Stryker, 2010; Keller et al., 2012), consistent with a preconditioning/priming to strengthen primary sensory processing. One expects similar examples of structural plasticity are likely in formation and processing of memories in the dentate gyrus and hippocampus (Clelland et al., 2009; Berron et al., 2016; van Dijk et al., 2023).

## Acknowledgements

We thank the OHSU Advanced Light Microscopy Core for imaging support. We acknowledge the use of viral vectors including AAV- hM3Dq (UNC Viral Core), AAV-flex-TeLC (gift from Peer Wulff), and retrograde AAV-hSyn-DIO-EGFP (Addgene #50457). We thank the OHSU Transgenic Core and the Mutant Mouse Resource and Research Center (MMRRC) for providing the Sim1-Cre mouse line, and Liqun Luo for facilitating access to the Fos-TRAP mice. Fos-TRAP and Sim1-Cre mice were essential to this study, and we are grateful for access to these transgenic resources. This work was supported by NS117371 (GLW) and the Dixon Professorship (GLW). The contents of this manuscript do not represent the views of the U.S. Department of Veterans Affairs or the United States Government (ES).

## Relative Contributions

Designed experiments - CC, GLW

Performed experiments and/or analyzed data, CC, AS, AV, AE

Assisted with experiments - MK, TM

Wrote and edited manuscript - CC, ES, GLW

## Data availability statement

All source data are available on reasonable request or at https://osf.io/6x7wj/overview?view_only=0049ef9e686d4e23839fba902221d1a5

## METHODS

### Mice

All procedures were performed according to the National Institutes of Health Guidelines for the Care and Use of Laboratory Animals and were in compliance with approved IACUC protocols at Oregon Health and Science University. Both female and male mice from a C57BL6/J background were used for experiments, aged 6–8 weeks at the time of surgery. TRAP mice were heterozygous for the FosCreERT2 allele (JAX#021882; (Guenthner et al., 2013); some were also heterozygous for the B6;129S6-Gt(ROSA)26Sortm14(CAG-tdTomato)/Hze/J(Ai14;JAX#007908) allele for experiments involving tdTomato labeling. The Sim1-Cre line, which expresses Cre under the control of the Single minded homolog-1 (Sim1) promoter, was generated by the OHSU transgenic core and obtained from MMRRC 034614-UCD (strain name: Tg(Sim1cre)KH21Gsat/Mmucd). Sim1-Cre mice were bred to be heterozygous for the Cre transgene by crossing a male Sim1-Cre mouse carrying the transgene with female C57BL6/J mice or with female homozygous for the B6;129S6-Gt(ROSA)26Sortm14(CAG-tdTomato)/Hze/J(Ai14;JAX#007908) allele for experiments involving tdTomato labeling.

### Viral Constructs

To express the Gq-coupled hM3D DREADD (designer receptor exclusively activated by designer drug) selectively in entorhinal cortex axons projecting to the MML or OML, we stereotaxically injected an AAV5 viral construct (1x10e13 particles/ml, University of North Carolina Viral Core) into the medial or lateral entorhinal cortex. Laminar-specific labeling was confirmed by examination of viral transduction specificity (via fluorescence microscopy for the appropriate fluorophore) in the entorhinal cortex and in the dentate gyrus molecular layer. For the DREADD experiments, injections that infected both the lateral and medial entorhinal cortex were excluded from the analysis. The pAAV-hSyn-DIO-EGFP (AAV Retrograde) was obtained from Addgene (50457-AAVrg). To selectively silence EC axons, we used a custom-made tetanus toxin light chain (TeLC) virus containing the GFP-tagged TeLC reading frame inverted in a flip-excision (FLEX) cassette (AAV-FLEX-TeLC-GFP), which was a kind gift by Peer Wulff (Murray et al., 2011). AAV-FLEX-GFP (also a gift from Dr. Wulff) was used as a control. We also used AAV5-TeLC-mCherry. Both constructs were packaged by Vectorbuilder with an AAV5 capsid, and injected at a titer of 1 x 10e13 particles/ml.

### Stereotaxic injections of AAV-based vectors

Mice were anesthetized using an isoflurane delivery system (Veterinary Anesthesia Systems Co.) by spontaneous respiration and placed in a Kopf stereotaxic frame. Skin was cleaned with betadine and topical lidocaine was applied before an incision was made. The dentate gyrus granule cell layer was targeted by placing Burr holes at −1.9 mm anteroposterior; ±1.1 mm lateromedial; −2.5 mm, −2.3 mm dorsoventral from Bregma. Using a 10 µl Hamilton syringe with a 30 ga needle and the Quintessential Stereotaxic Injector (Stoelting), 1 µl viral stock was delivered at each site at 0.25 µl/min. The syringe was left in place for 1 min after each injection before slow withdrawal. Injections into the lateral entorhinal cortex were made at the following coordinates: anteroposterior (from bregma), −3.4 mm; lateromedial (from bregma), ±4.0 mm; dorsoventral (from brain surface), −2.4 mm. Injections into the medial entorhinal cortex were made at the following coordinates: anteroposterior (from bregma), −4.5 mm; lateromedial (from bregma), ±3.0 mm; dorsoventral (from brain surface), −3.2 mm. Stereotactic coordinates for the middle molecular layer of the dentate gyrus were as follows: anteroposterior (from bregma), −2.0 mm; lateromedial (from bregma), ±1.2 mm; dorsoventral (from brain surface), −2.0 mm. Stereotactic coordinates for the outer molecular layer of the dentate gyrus were as follows: anteroposterior (from bregma), −2.0 mm; lateromedial (from bregma), ±1.2 mm; dorsoventral (from brain surface), −1.8 mm. Injections in both OML/MML regions were performed using the same procedure described above, with 1 µl of viral stock delivered at each site at 0.25 µl/min. The syringe was left in place for 1 min after each injection before slow withdrawal to ensure proper diffusion of the viral solution. To control for differences in the degree of innervation from the medial and lateral entorhinal cortex (Dolorfo and Amaral, 1998; Ohara et al., 2013), injections in the dentate gyrus were targeted to the intermediate region across the septotemporal axis (van Groen et al., 2003; Strange et al., 2014), which receives approximately equal innervation from the medial and lateral entorhinal cortex. The skin above the injection site was closed using veterinary glue. Animals received post-operative analgesia with topical lidocaine and flavored acetaminophen in the drinking water.

### TRAP induction

Tamoxifen was dissolved at 20 mg/ml in corn oil by sonication at 37°C for 5 min. The dissolved tamoxifen was then stored in aliquots at –20°C for up to several weeks or used immediately. The dissolved solution was injected intraperitoneally at 150 mg/kg. For exercise-TRAP, tamoxifen was administered 23 hr before exposure to the running wheel. Experiments were begun as early as 48 hr after tamoxifen administration to allow for reporter expression for the experimental timepoints at 3, 5, or 7 days post-tamoxifen (see Figure legends).

### Voluntary exercise assay and enriched environment design

Both home cage and exercise groups were housed together until 7 days before the experiment, after which they were singled housed in oversized (rat) sedentary cages (43 × 21.5 cm^2^) to allow acclimation to the novel environment before tamoxifen administration. At 23 hr post-tamoxifen injection, a running wheel was introduced in the cage of animals in the exercise group and mice had free access to running or exploring for 2 hr at the beginning of the dark period, after which the running wheel was removed from the cage. Total distance (km) was measured using an odometer. All mouse groups were handled for 5 days before tamoxifen administration. For enriched environment experiments, mice were housed in standard rat cage equipped with toys, houses, and a maze-like tube system for 6 or 24 hrs. then assessed for c-Fos immunohistochemistry. A separate cohort of mice were acclimated to an open field arena on day 1 and then returned on day 2 to assess total distance traveled with or without the EE objects. Mice were placed individually in an open field arena (40 × 40 cm²) with high walls to prevent escape. The arena floor was divided into a grid, and movements were recorded for 20 minutes. Total distance traveled was analyzed.

### Immunohistochemistry

Mice were terminally anesthetized, transcardially perfused with saline and 20 ml of 4% paraformaldehyde (PFA), and brains were post-fixed overnight. Coronal sections (100 μm) of the hippocampus were collected and permeabilized in 0.4% Triton in PBS (PBST) for 30 min. Sections were then blocked for 30 min with 5% horse serum in PBST and incubated overnight (4°C) with primary antibody in 5% horse serum/PBST. After extensive washing, sections were incubated with the appropriate secondary antibody conjugated with Alexa 488, 568 or 647 (Molecular Probes), for 2hrs at room temperature. They were then washed in PBST (2 × 10 min) and mounted with Dapi Fluoromount-G (SouthernBiotech). The primary antibodies used were: anti-c- fos (1:500, Santa-Cruz), anti-tdTomato (1:500, Clontech), anti-VGLUT2 (1:500, Synaptic Systems), anti-reelin (MBL, 1:200) and rabbit anti-calbindin D-28K (SWANT, 1:2500), anti-doublecortin (DCX, 1:500, Millipore, Billerica, MA), anti-BrdU (1:500, Abcam). All antibodies have been well characterized in prior studies in our laboratory, and staining was not observed when the primary antibody was omitted.

### Imaging and morphological analysis

For imaging we used an LSM 900 laser scanning microscope (Carl Zeiss MicroImaging; Thornwood, NY). All cell and dendritic spine counting were done manually with ImageJ and Imaris (National Institutes of Health; Bethesda, MD). For quantification of immunopositive cells, six hippocampal slices at set intervals from dorsal to ventral were stained per animal; 3–6 animals were analyzed per group. A 49 μm z-stack (consisting of seven optical sections of 7 μm thickness) was obtained from every slice at 20x objective. Positive cells were counted per field from every z-stack, averaged per mouse and the results were pooled to generate group mean values. For analysis of spine density of dentate granule cells, sections were imaged with a 63x objective with 2.5x optical zoom. The span of z-stack was tailored to the thickness of a single segment of dendrite with 0.1 µm distance between planes. Dendritic spines were imaged and analyzed in the middle (MML) and outer molecular (OML) layers of the dentate gyrus. The MML and OML were distinguished based on the pattern of VGluT2 immunofluorescence, which begins at the border between the inner molecular layer and middle molecular layer. MML dendritic segments were therefore imaged at the beginning of the VGluT2 staining nearest the granule cell body layer, whereas OML dendritic segments were imaged at the distal tip of the molecular layer. The same microscope settings (laser intensity and gain) were used for each experimental group analyzed. Slides were coded, and slides were imaged/analyzed by an investigator blinded to experimental condition.

### Brain slice physiology

To validate the TeLC-GFP construct, adult mice were used 3-4 weeks following AAV-FLEX-TeLC-GFP injection. For slice preparation, animals were deeply anesthetized using isoflurane followed by an IP injection of 0.8mL 2% Avertin (Tribromoethanol, Sigma). They were transcardially perfused with an ice-cold choline chloride solution (containing in mM: 110 choline chloride, 1.25 NaH_2_PO_4_, 2.5 KCl, 1.3 Na-ascorbate, 7 MgCl_2_, 1 CaCl_2_, 25 NaHCO_3_, 10 D-Glucose) which was adjusted to an osmolarity of 310 mOsm and saturated with 95% O_2_ – 5% CO_2_ gas mixture. 300 <u>μ</u>m thick coronal slices were cut in ice-cold choline solution using a vibrating microtome (VT1200, Leica Microsystems). One hemisphere was marked with a notch prior to slicing to allow for differentiation of the injected side. Slices were then incubated at 34 degrees in artificial cerebral spinal fluid (ACSF) for 30 minutes, and allowed to sit for an additional 30 minutes at room temperature before recording. Slices were then placed in a recording chamber and perfused with oxygenated, room temp ACSF at ∼2.5 mL/min. Extracellular field potential recordings were obtained via a 1MOhm borosilicate glass pipette filled with 3M NaCl solution placed 250-350um away from a bipolar stimulating electrode in either the MPP or the LPP. To ensure correct placement of stimulating electrodes, the recording electrode was placed in the opposite pathway of interest to validate an expected reversal of the field potential polarity. Slices were stimulated at 50% of maximum response (between 4 and 6V, typically; 100 µsec), except for slices that were silenced, which the stimulation was increased to 11V to ensure no response was elicited. All signals were amplified by a MultiClamp 700B (Molecular Devices, San Jose, CA) amplifier, low pass filtered at 6kHz and sampled at 5kHz. using a National Instruments analog-to-digital board (NiDAQ) and collection by IgorPro-based software (WaveMetrics, Lake Oswego, OR, USA). For analysis, multiple sweeps from the same recording were averaged to calculate amplitudes. Paired pulse ratios (PPR) were calculated by comparing the maximum rising slope of each field response in the MPP and LPP. Max slope was determined between points on the ∼25ms following the afferent volley prior to the field EPSP peak. Calculations were done with IgorPro or in Excel.

### BrdU treatment

To birthdate newborn cells in the adult hippocampus, mice were injected intraperitoneally with Bromodeoxyuridine (BrdU, Sigma-Aldrich, St. Louis, MO) at 300 mg/kg twice with a 4 hr interval between doses on a single day, and sacrificed at either 21 or 28 days after injection. This pulse-chase protocol was chosen to saturate mitotic cell labeling within a single cell cycle as determined previously (Cameron and McKay, 2001).

### Statistical analysis

Sample sizes were based on pilot experiments with an effect size of 20% and a power of 0.8. Littermates were randomly assigned to control or exercise groups. Criteria were established in advance based on pilot studies for issues including data inclusion, outliers, and selection of endpoints. Criteria for excluding animals from analysis are listed in the methods. Mean ± SD was used to report statistics for all experiments, apart from electrophysiology experiments where mean ± SE was used. The choice of statistical test, test for normality, definition of N, and multiple hypothesis correction where appropriate are described in the figure legends. Unless otherwise stated, all statistical tests were two-sided. Significance was defined as p<0.05. All statistical analyses were performed in Prism.

## Key Resources Table

| Reagent type | Designation | Source or reference | Catalog Number | RRID |
| --- | --- | --- | --- | --- |
| Genetic reagent (mouse) | c-FOScreERT2 | Guenthner et al., 2013 | N/A | N/A |
| Genetic reagent (mouse) | Ai14 (B6;129S6-Gt(ROSA)26Sortm14(CAG-tdTomato)/Hze/J) | The Jackson Laboratory | 007908 | RRID:IMSR_JAX:007908 |
| Genetic reagent (mouse) | Sim1:Cre | Generated at OHSU, obtained from MMRRC | Tg(Sim1cre)KH21Gsat | RRID:MMRRC_036051-UCD |
| Recombinant DNA reagent | pAAV-hSyn-DIO-EGFP | Addgene | 50457 | RRID:Addgene_50457 |
| Recombinant DNA reagent | AAV-FLEX-TeLC | Gift from Peer Wulff | N/A | N/A |
| Antibody | anti-c-Fos (rabbit polyclonal) | Santa Cruz Biotechnology | sc-52 | RRID:AB_2106783 |
| Antibody | anti-tdTomato (rabbit polyclonal) | Clontech | 632496 | RRID:AB_10013483 |
| Antibody | anti-VGLUT2 (rabbit polyclonal) | Synaptic Systems | 135402 | RRID:AB_887883 |
| Antibody | anti-reelin (mouse monoclonal) | MBL | D223-3 | RRID:AB_592179 |
| Antibody | anti-calbindin D-28K (rabbit polyclonal) | Swant | CB38a | RRID:AB_10000340 |
| Antibody | anti-doublecortin (DCX, goat polyclonal) | Millipore | AB2253 | RRID:AB_1586992 |
| Antibody | anti-BrdU (rat monoclonal) | Abcam | ab6326 | RRID:AB_305426 |
| Chemical compound, drug | Tamoxifen | Sigma-Aldrich | T5648 | N/A |
| Chemical compound, drug | Bromodeoxyuridine (BrdU) | Sigma-Aldrich | B5002 | N/A |
| Software, algorithm | ImageJ | NIH | N/A | RRID:SCR_003070 |
| Software, algorithm | Imaris | Bitplane | N/A | RRID:SCR_007370 |
| Software,<br>algorithm | Igor Pro 8 | WaveMetrics | N/A | RRID:SCR_000325 |

## FIGURE LEGENDS

**Figure Supplemental 1. Exercise-TRAPed granule cells are mature granule cells.**

A. Representative images of exercise-TRAPed granule cells (red) co-stained for the mature granule cell marker calbindin (Cb) and the immature granule cell marker doublecortin (DCX) after 2hr of voluntary exercise, indicated that the exercise-TRAPed cells were mature (Cb^+^) granule cells. (Cb⁺/exercise-TRAPed⁺ overlap: 72.4 ± 5.2%; DCX⁺/ exercise-TRAPed overlap: 1.2% ± 0.01%, n=5, p < 0.0001). Scale bar = 100 µm.

B. Experimental design for BrdU birth-labelling of exercise-TRAPed granule cells. Two-month-old male and female Fos-TRAP:TdTomato mice received two injections of BrdU per day (200mg/kg, i.p.). Mice were injected once with tamoxifen (150 mg/kg) 24 hr before exposure to 2 hr of voluntary exercise at 21 or 28 days post-BrdU. Mice were sacrificed 2 days after exposure to the running wheel. (Middle) Representative image of the dentate gyrus of Fos-TRAP:TdTomato mice 3 weeks after BrdU injection: BrdU (green), Exercise-TRAPed cells (red). See text for quantification. Scale bar = 200 µm.

