## Supplementary figures and images for "Biased synaptic activation of dentate granule cells by exercise reflects inputs from the lateral entorhinal cortex"

### Figure S1

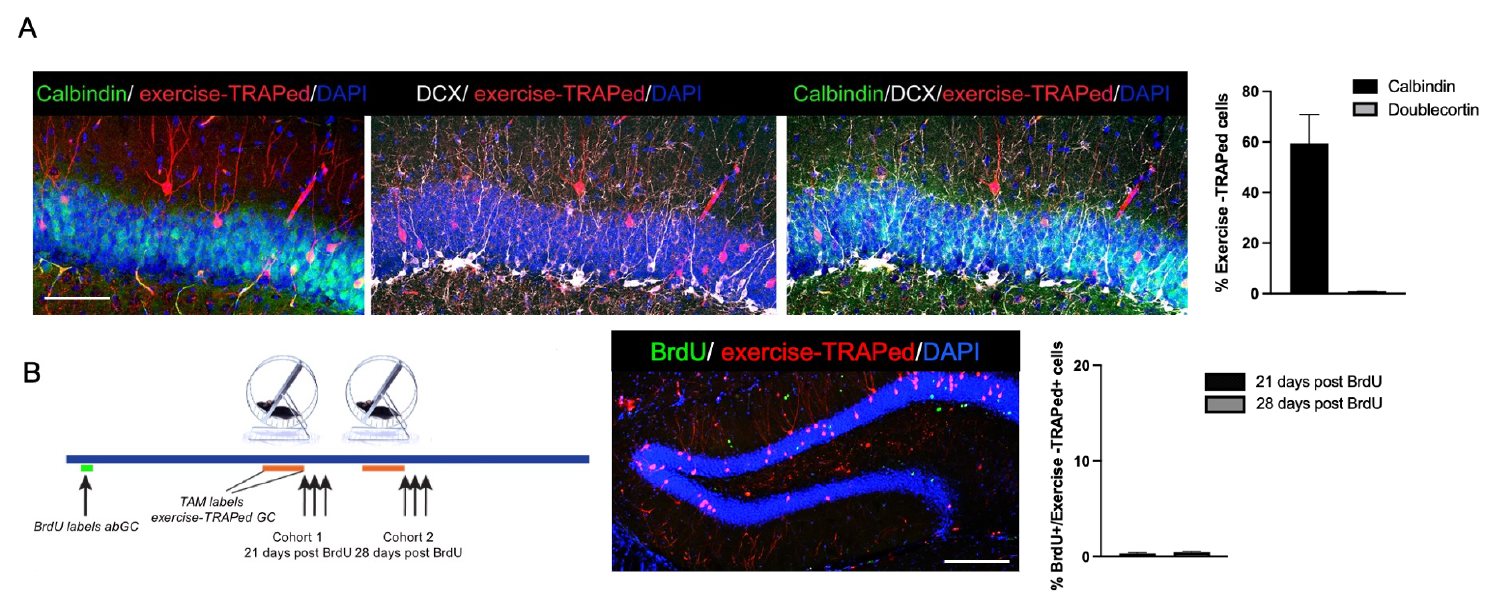
